# Heterotopic reduction of forelimb progenitors underpins development of the vestigial emu wing; implications for vertebrate limb evolution

**DOI:** 10.1101/2022.11.23.516993

**Authors:** Axel H Newton, Sarah M Williams, Belinda Phipson, Andrew J Pask, Andrew T Major, Craig A Smith

## Abstract

The tetrapod limb has evolved a broad diversity of form and function, enabling vertebrates to adapt to a wide range of environments. One of the most striking examples of this is the evolution of powered flight in birds. However, during the avian radiation, flightless ratites (Paleognathae) secondarily evolved multiple instances of wing reductions, utilizing distinct molecular mechanisms, and displaying heterochrony with flighted birds (Neoaves). Among ratites, the emu displays the most marked reduction of wing anatomy, developing a small wing with a single clawed digit. This major divergence from the typical avian pattern provides a robust model to determine the mechanisms underpinning developmental plasticity in the tetrapod limb. To interrogate the ontogenetic origins of vestigial wing development, we applied comparative single cell transcriptomics of the embryonic emu forelimb field with the chicken, as it possess a typical wing. Stunted wing development in the emu occurs through a heterotopic reduction of lateral plate mesoderm (LPM)-derived limb progenitor cells, and accompanying downregulation of genes associated with LPM development and early limb initiation. Here, forelimb bud initiation and outgrowth is altered by changes to the specification and allocation of limb progenitor cells, acting as an underlying template for pre- and post-patterning mechanisms. These findings provide novel insights into the evolution of limb development in tetrapods, and emphasizes an important, yet underappreciated role for heterotopy as major driver of limb disparity.

**Significance statement:** Animals exhibit a striking variety of morphologies, yet our understanding of how these forms evolve and arise during development are lagging. The emu is a large, flightless bird which has evolved a vestigial wing. This model allows detailed interrogations into the mechanisms which underlie evolution and development of a distinct morphological structure. Through application of comparative single cell transcriptomics between the emu and chicken, we identify an early embryological mechanism, heterotopy, underlying the evolution and vestigial development of the emu wing. This mechanism features reduced allocation of limb bud progenitor cells in the lateral plate mesoderm (LPM). This data provides new information on the mechanisms that underpin evolutionary diversity in morphological features.

## Introduction

One of the most striking modifications of the tetrapod limb was evolution of the avian wing to achieve powered flight. This transition can be traced across archosaur evolution ^1^, occurring as a reduction from five to three digits, evolution of flight feathers and modification of the limb musculature ^2,3^. However, some birds have secondarily lost the ability to fly, accompanied by significant morphological changes to wing structure and typically vestigial development. The Paleognathae, or ratites, are a clade of flightless birds, which include the ostrich, emu, rhea, cassowary, and kiwi. Despite their shared evolution towards terrestrialism, phylogenomics have revealed that ratites are paraphyletic, with each lineage losing the ability to fly independently, except for the tinamou which has retained flight ^4–7^. Studies into the genomic basis of this reduction has revealed ratites possess homoplasy within cis-regulatory elements surrounding genes associated with limb development ^8^, suggesting selection has convergently targeted shared loci during the evolution of flight loss. However, other evidence suggests vestigial wing development amongst ratites occurs through distinct developmental processes ^9^. As such, the precise mechanisms which underlie development of the vestigial ratite forelimb are largely unknown, especially during early limb specification and outgrowth. Owing to their ease of experimental manipulation *in ovo*, the ratite forelimb provides a compelling model for comparative interrogations into the molecular foundations of limb development and disparity, particularly when compared to flighted birds.

The limbs arise through a series of embryological events that begin in the LPM. During embryogenesis, the undifferentiated LPM undergoes dorsoventral subdivision along the anteroposterior axis of the embryo into the somatic and splanchnic LPM cell layers, producing the embryonic coelom ^10,11^. In the first stages of limb development, colinear activation of HOX genes along the anterior-posterior axis define the presumptive fore and hindlimb fields within the established somatic LPM ^12–15^. BMP signals from the overlying ectoderm specify somatic LPM progenitors towards a limb fate, observed by expression of *PRRX1, IRX3* and *TWIST1* ^11,16,17^. Epithelial cells in the somatic LPM monolayer then undergo EMT to produce the limb bud mesenchyme ^16,18,19^, which in the forelimb, is dependent on the limb initiation factor *TBX5* ^14,20–23^. TBX5 activates *FGF10* in the limb bud mesenchyme, which establishes a positive feedback loop with *FGF8* in the overlying ectoderm to stimulate outgrowth of the limb buds ^21,24–26^. Disruption of early ectodermal BMP signalling ^16^, *Tbx5* during the window of limb initiation, or later activation of *Fgf10* or *Fgf8,* all result in a failure of forelimb development ^22,27,28^. As such, the evolution of divergent limb morphologies across vertebrates, including bat wings, mammalian digit reductions, dolphin fin and jerboa hindlimb expansions ^29–34^, are thought to occur via post-patterning of the established limb primordia ^34,35^.

Of the flightless ratites, the emu (*Dromaius novaehollandiae*) is an experimentally tractable model which possesses the most dramatically reduced forelimb, characterized by a single clawed digit ^36,37^. Studies have begun to examine the underlying mechanisms contributing to its delayed and reduced forelimb, with several events associated with its stunted outgrowth and differentiation (reviewed in ^35^). Relative to the chicken, which displays a typical wing, the emu shows delayed differentiation of the somatic LPM ^38^; a reduced *TBX5-*positive forelimb field ^39^; attenuated activation of *FGF10* ^38^ and delayed expression of *SHH* ^40^ in the early wing bud mesenchyme; and co-option of the cardiac-specific transcription factor *NKX2.5* into the limb myogenic precursors, though to repress limb outgrowth ^41^. Interestingly, while each of these mechanisms occur in a sequential cascade, they appear to stem from an initial delay in formation of the emu’s forelimb domain from the LPM ^38,39^. This suggests that the molecular basis of differential wing morphogenesis may arise earlier during embryonic development than currently appreciated, during the initial formation of the somatic LPM.

To define the ontogenetic and molecular basis of reduced wing development in the emu, we applied single cell transcriptomics during early limb specification and initiation, drawing evolutionary comparisons with our recent work in the chicken ^16^. Through stage-matched sampling of cells in the developing forelimb field during LPM specification, somatic LPM differentiation, limb initiation and early outgrowth, we resolve the ambiguity in previous published studies. We demonstrate that the emu does not possess heterochronic limb development as previously thought ^38,40,42,43^, but rather has evolved a heterotopic reduction of limb progenitors from within the somatic LPM, preceding activation of the canonical limb development pathway. Where heterochrony describes changes to rate or timing of development, heterotopy is a lesser studied phenomenon referring to changes in spatial arrangement or quantities in an embryo ^44^. The reduced specification of forelimb progenitors observed in the emu are further accompanied by reduced and delayed expression of known LPM and limb genes, suggesting the reduced commitment of forelimb progenitors is encoded during patterning of the early LPM. These findings provide new perspectives into our understanding of limb development and evolution, where heterotopic allocation of progenitors during embryonic development can influence the divergent development of different limb morphologies between species.

## Results

We utilized the 10x Chromium system to generate single-cell transcriptomic profiling of cells within the developing chicken and emu forelimb field. The overall embryology of the emu closely resembles flighted birds, including that of the chicken ^12,42,45^, with forelimb initiation occurring at roughly equivalent developmental stages ^38^. Stage-matched embryonic forelimb field tissues of chicken embryonic day e1.5, e2.5 and e3.5 ^16^, and emu e3.5, e4.5 and e5.5– corresponding to Hamburger-Hamilton stages HH10, HH14 and HH18, respectively (Figure 1a), were dissociated and subject to single cell RNA sequencing. After quality filtering, we identified a total of 15355 chicken cells and 16748 genes, and 16149 emu cells with 15628 genes. To remove low read count or low diversity cells, we applied a strict filtering threshold for cells with high mitochondrial counts and ≥2000 UMI per cell. This produced a final dataset of 3262 high quality cells in chicken, and 2678 cells in emu (summarized in Table S1) used in the analysis. Cell transcriptomic relationships were visualized with global t-distributed Stochastic Neighbour Embedding (tSNE) dimension reduction, which produced corresponding cell type distributions within the chicken and emu presumptive forelimb field datasets, displaying clear separation of clusters comprising the embryonic ectoderm, mesoderm, endoderm and vasculature (Figure 1b-d). Independent unsupervised clustering of the chicken and emu HH10, HH14 and HH18 presumptive forelimb datasets revealed 13 transcriptionally distinct cell clusters (c) in the chicken (Figure 1c) as previously determined ^16^, and 16 clusters in the emu (Figure 1d). Tissue and cell type identities were assigned to each cluster based on their differential gene expression profiles (observed as differences in log-fold change between clusters; Figure 1e,f) and corresponding spatiotemporal expression patterns observed throughout the developing chicken embryo (*Gallus Expression In situ Hybridization Analysis*: GEISHA ^46,47^). Annotated cell types were largely in agreement between the two species datasets showing single clusters representing the endoderm, vasculature and RBCs, two-three clusters representing the ectoderm, and several clusters for the mesoderm, including paraxial, intermediate, and lateral plate derivatives (Figure 1c-d).

**Figure 1.**
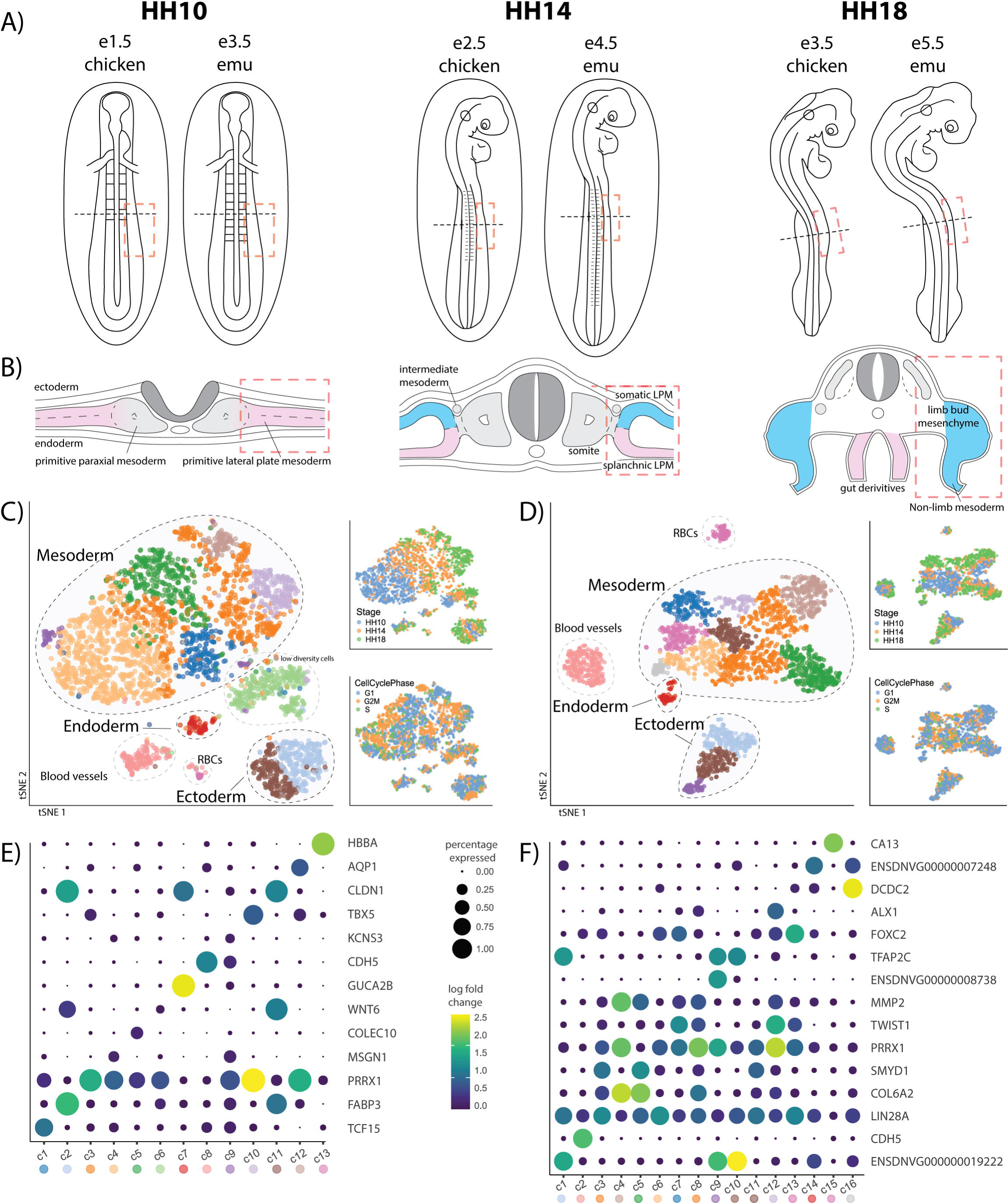
Identification of cell types in the chicken and emu forelimb field. (A) Cells were isolated from chicken embryonic day - E1.5, E2.5 and E3.5, and emu E3.5, E4.5 and E5.5 to sample all major tissues in the developing forelimb field. (C,D) tSNE visualisation separated chicken and emu cells based on stage and germ layer origin (B). (E,F) Unique gene expression profiles were detected for each major cluster of chicken and emu cells, which were largely in agreement, recapitulating shared cell clusters with overlapping marker genes.

### The emu exhibits reduced commitment of forelimb progenitors from the somatic LPM

To identify the cellular basis of differential emu wing bud development, the chicken and emu datasets were integrated into a single merged atlas. A common set of genes with 1:1 orthology between both species were identified and used to subset the datasets to only contain common gene IDs (n=10415). The datasets were then merged and visualized through tSNE dimension reduction, which revealed batch effects between species (Figure S1a). These were removed through mutual nearest neighbours (MNN) correction ^48^, producing a combined, homogeneous chicken-emu dataset with equivalent clusters corresponding to embryonic ectoderm, mesoderm, endoderm and vasculature (Figure S1b).

As the limb mesenchyme arises from the embryonic mesoderm, the merged chicken-emu dataset was further subset into cells of mesodermal origin (Figure 2a, Figure S2a). Additional unsupervised clustering of the merged dataset revealed 9 sub-clusters, corresponding to all know tissue types within the limb field mesoderm. Namely, these annotated cell types consisted of early (undifferentiated) mesoderm, paraxial and intermediate mesoderm, undifferentiated lateral plate mesoderm (LPM), somatic and splanchnic LPM, visceral (gut) precursors, limb bud mesenchyme and extra-embryonic / flank LPM (Figure 2b,c), recapitulating annotated cell types within the developing chicken limb (Figure S2c) ^16^. Of interest, *NKX2-5* was not detected in chicken cells, but was active in emu paraxial mesoderm (somites) and somatic LPM prior to formation of the limb bud mesenchyme (Figure S2b), supporting previous observations ^38,41^. Cell lineage trajectories across the merged dataset were determined with monocle3 ^49^, supporting the subdivision of the undifferentiated LPM into the splanchnic/gut and somatic/limb lineages (Figure 2b), corresponding with clusters and trajectories previous determined in the chicken (Figure S2c) ^16^. Despite attempted stage-matched sampling of chicken and emu tissues based on developmental series’ ^42,45^, initial examination of stage-matched cell types revealed that the emu sampling largely failed to recover early undifferentiated mesoderm present in the chicken (Figure 2a-c). Instead, emu embryonic day (E)3.5 cells more closely resembled HH14 (Figure 2e). By E5.5 (∼HH18) however, emu cell types matched those of HH18 chicken samples (Figure 2c-d), suggesting our three emu samples were equivalent to chicken stages ∼HH12 to HH18. As our emu sampling sufficiently captured the transition from undifferentiated LPM, through its subdivision into somatic / splanchnic LPM and into the limb mesenchyme and visceral cell fates, respectively, we were able to examine cell type proportions during limb initiation.

**Figure 2.**
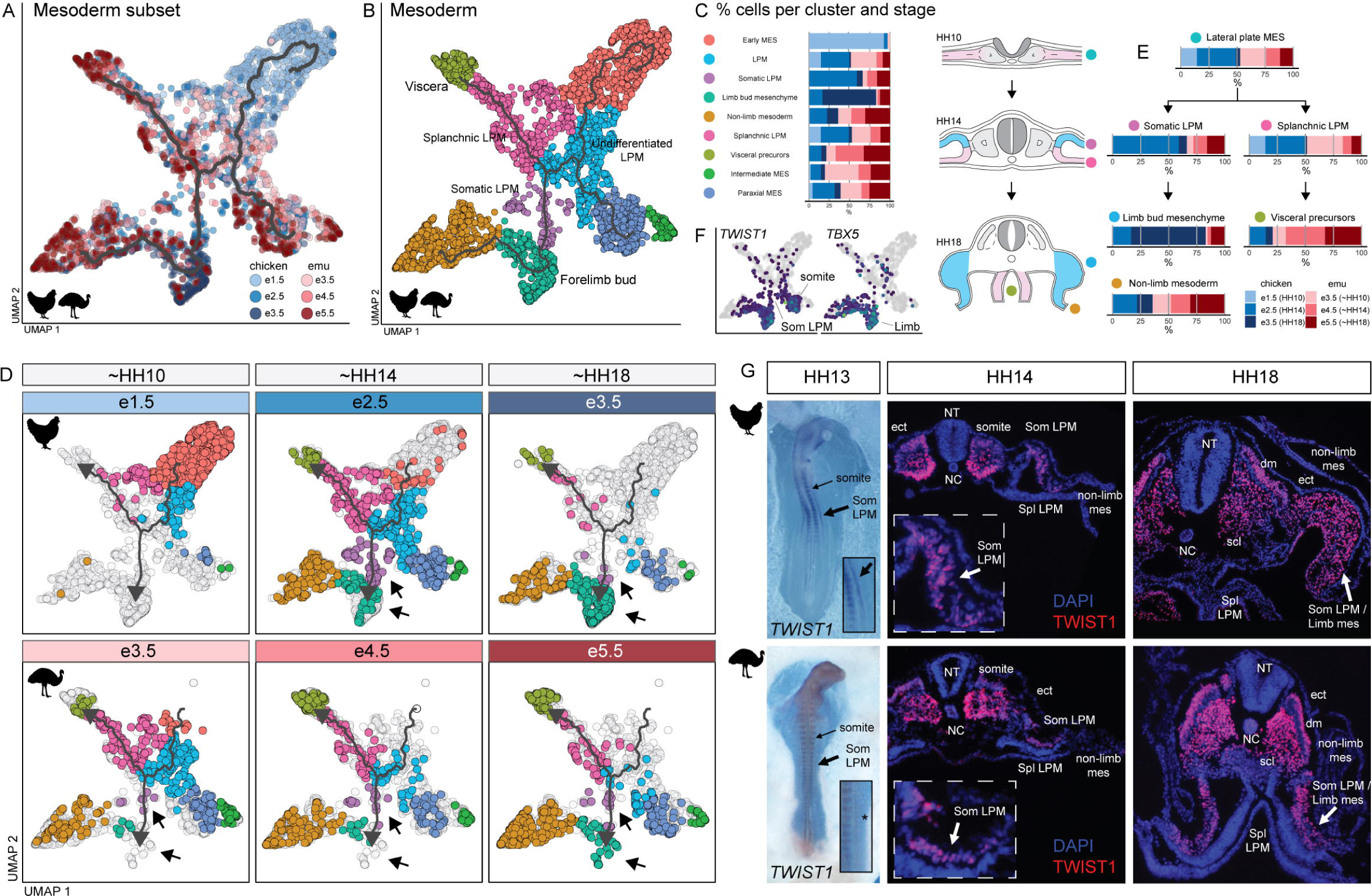
Comparisons of cell type proportions in the chicken and emu developing forelimb field. (A) The complete chicken and emu datasets were integrated to produce a unified atlas of the developing forelimb field, recapitulating tissues from all germ layers. (B) Mesodermal cells were subset and clustered to produce (C) 9 clusters covering known mesodermal cell types. Total proportions of cells were identified for each cluster, corresponding to each species and stage. (D) A trajectory was fit from the undifferentiated LPM to the limb bud mesenchyme (B) and the presence and numbers of cells for cluster, per species per stage were identified. This revealed a reduced allocation of somatic LPM cells in the emu from undifferentiated LPM precurosrs (arrow), relative to the splanchnic LPM and to the chicken (E). Immunofluorescent labelling of the somatic LPM marker TWIST1 corroborated the reduced proportions of emu somatic LPM and limb bud cells, showing an overall reduced number of TWIST1+ cells in the emu, relative to the splanchnopleure and the chicken at roughly equivalent stages. som = somatopleure/somatic LPM; spl = splanchnopleure/splanchnic LPM; dm = dermomyotome; scl = sclerotome; nc = notochord; nt = neural tube.

To determine whether the emu indeed possesses delayed differentiation of the somatic LPM compared with the chicken ^38^, we investigated differences in cell type proportions between the chicken and emu during LPM subdivision, somatic LPM differentiation and formation of the limb mesenchyme. The LPM undergoes dorsoventral subdivision into the somatic and splanchnic layers in equal ratios via transient formation of epithelial rosettes ^10^, producing the coelom along the anteroposterior axis of the embryo ^11^. Prior to subdivision, we identified approximately equal numbers of chicken and emu undifferentiated LPM cells in our datasets (Figure 2c-e). After LPM subdivision, the overall proportions and numbers of somatic and splanchnic cells between the chicken and emu were roughly maintained, though with the emu displaying a reduction in the number of somatic LPM cells (Figure 2d-e) compared to chicken. However, during and after limb initiation, the emu showed a significant reduction in limb bud mesenchyme cells, and increase in non-limb mesoderm and splanchnic-derived cells, relative to that of the chicken (Figure 2c-e).

To visualize these observations *in vivo*, we performed immunofluorescent labelling of the limb-progenitor and limb mesenchyme fate marker TWIST1 ^16^ (Figure 2f) in chicken and emu embryos. Whole mount *in situ* hybridization of *TWIST1* in chicken and emu embryos revealed a smaller, narrower and less intense band of somatic-LPM derived forelimb progenitors in the developing emu forelimb field (Figure 2g). This was further explored in HH14 and 18 forelimb tissue sections, before and after limb initiation. Immunostaining for TWIST1 further revealed that the emu possesses a smaller overall field of TWIST1^+^ somatic LPM-derived limb progenitor cells during limb initiation, compared to relative abundances of TWIST1+ somites and splanchnic LPM cells observed in the chicken (Figure 2g). By HH18, the chicken had a distinct limb bud full of proliferative limb bud mesenchyme, whilst the emu showed attenuated pools of limb bud progenitors. Together, these results emphasize that specification of the emu somatic LPM is not delayed ^38^, rather there is a reduced commitment of somatic LPM-derived limb-progenitors within the presumptive forelimb field, prior to limb initiation.

#### The emu shows differential gene expression dynamics underlying wing heterotopy

To understand the molecular basis of the reduced specification of the emu somatic LPM, we investigated differentially expressed genes in the emu LPM, somatic LPM and forelimb bud mesenchyme. This was first analysed through pseudobulk differential expression (DE) analysis of emu cell clusters with the corresponding clusters in the chicken. This revealed differentially expressed emu genes in LPM tissues with known roles in various aspects of LPM development (Figure 3a). Of the differentially expressed genes within the undifferentiated LPM cluster, we identified emu-specific downregulation of the limb fate marker gene *PRRX1* ^50,51^ and increased expression of splanchnopleure marker *OSR2* ^52^, suggesting a reduced capacity for somatic LPM commitment. DE analysis of the somatic LPM clusters produced few statistically significant gene expression profiles, likely owing to low numbers of contributing cells in the emu dataset. However, in limb bud mesenchyme several differentially expressed genes were noted between chicken and emu, including emu-specific downregulation of limb-developmental genes *PRRX2, HOXD9*, *FGFR1* as well as *FGF10* which has previously been identified as a contributor to reduced emu limb outgrowth^38^. This confirms that while several limb bud genes are downregulated in the developing emu limb, somatic LPM fate genes are downregulated in the emu LPM prior to somatic LPM differentiation and limb bud formation.

**Figure 3.**
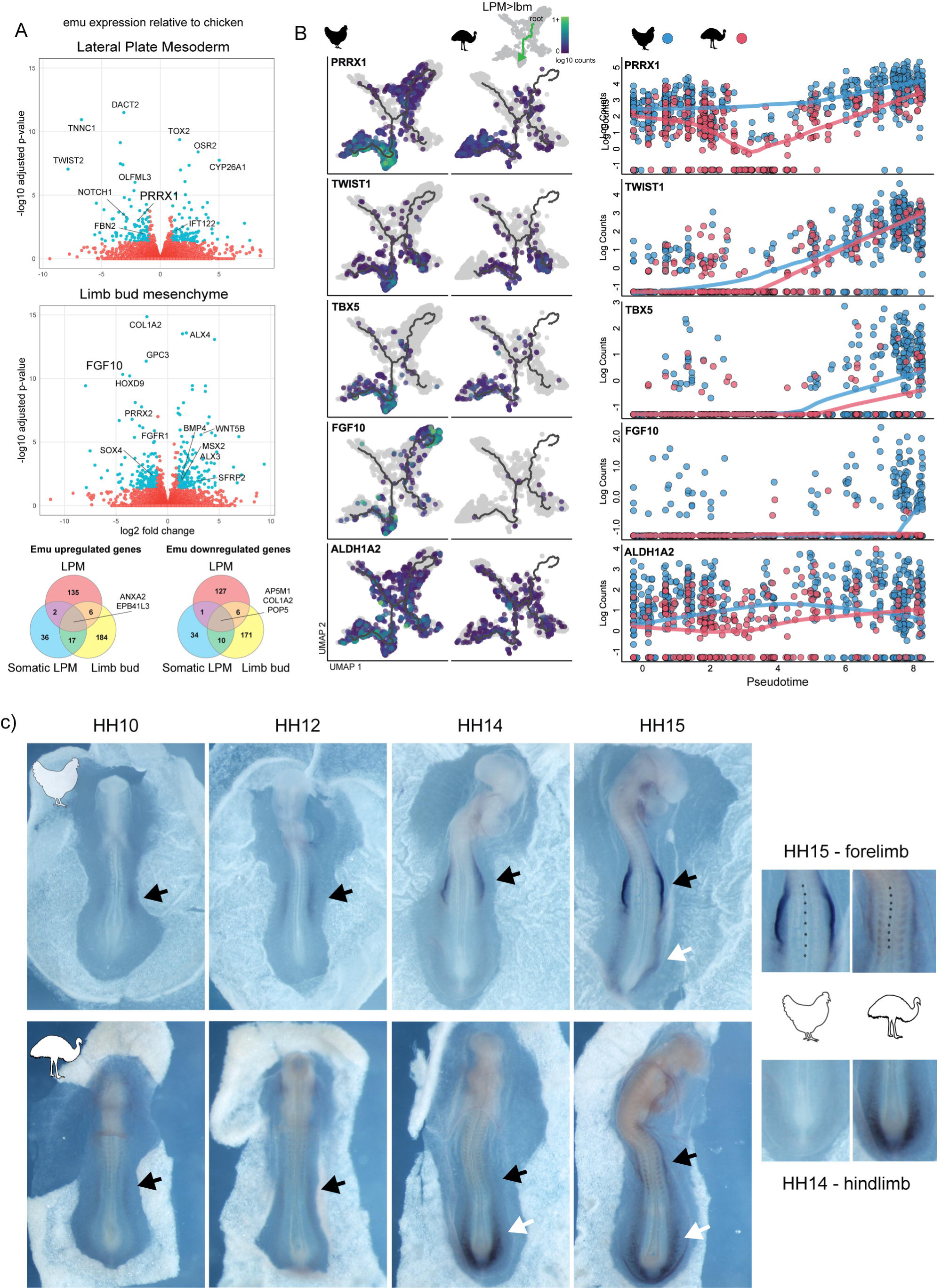
Gene expression dynamics underlying emu wing heterochrony. (A) Pseudobulk differential expression of emu lateral plate mesoderm and forelimb bud mesenchyme, relative to the chicken, revealed downregulation of several LPM markers (i.e. *PRRX1*) prior to limb bud formation, and limb genes (i.e. *FGF10*) post limb-initiation. (B) Visualization of differential expression of genes across pseudotime revealed downregulation of LPM and limb markers in the emu relative to the chicken across development. *PRRX1, TWIST1, TBX5*, *FGF10* and *ALDH1A2* each showed stage-specific decreases in expression, relative to chicken. (C) Spatial and temporal *in situ* visualization of *PRRX1* gene expression during chicken and emu forelimb development. This emu possessed reduced and delayed activation of PRRX1 in the developing forelimb field (black arrow) despite covering a similar number of somites, relative to the chicken. In contrast, *PRRX1* expression was increased and accelerated in the emu hindlimb relative to the chicken (white arrow). Together these suggest the emu has delayed and accelerated LPM specification in the forelimb and hindlimb, respectively.

To complement this static pseudobulk approach, we further investigated dynamic changes in gene expression during LPM and limb bud development across pseudotime. The root of the trajectory was set in the undifferentiated mesoderm/LPM boundary, where emu undifferentiated LPM cells were first detected (Figure 2, 3b). Cells were ordered along the LPM to limb mesenchyme lineage trajectory (Figure 2b, 3b) and pseudotime was calculated. We then examined dynamic gene expression changes across pseudotime, identifying genes which were differentially expressed between the chicken and emu along the trajectory. After excluding ribosomal genes, this revealed 1199 significantly downregulated, and 1566 significantly upregulated genes in the emu compared with the chicken during the LPM to limb transition. Functional gene ontology profiling of emu downregulated genes using g:GOSt showed enrichment of terms associated with embryonic limb morphogenesis (Figure S3). Of the top downregulated genes, these included known markers of LPM and limb bud development *PRRX1, PRRX2, TWIST1, WNT5A, SALL1, HOXD9, GLI3* and *FGF10*; and RA synthesis/response genes: *ALDH1A2, STRA6, RDH10, RARB* and *CYP26B*1 (Table S4). These gene lists complemented the pseudobulk results, but also revealed additional genes that were not captured by the static analysis.

Visualization of selected marker gene plots across pseudotime confirmed dynamic chicken and emu expression differences during LPM and limb development. Notably, we observed a delayed onset of *TWIST1* and sustained downregulation of somatic LPM markers *PRRX1* and *ALDH1A2* during emu LPM development (Figure 3b), as well as decreased activation and expression of limb initiation markers *TBX5* (Figure S4) and *FGF10,* confirming previous observations ^38,39^. As *PRRX1* was found to be downregulated in both static and dynamic DE analyses, and is a diagnostic lineage marker of the somatic LPM ^50,51,53,54^, we examined it’s spatiotemporal expression profile in emu and chicken embryos. Whole mount *in situ* hybridisation of *PRRX1* confirmed reduced expression in the developing emu forelimb field, appearing in a smaller domain of decreased intensity to that of the chicken (Figure 3c) showing similar patterns to *TWIST1* (Figure 2g). Curiously, this additionally highlighted *PRRX1* expression in the emu hindlimb was enhanced compared to the chicken, showing heterochronic acceleration and raising the possibility that allocation of somatic LPM progenitors of the hindlimb may be increased.

To test whether *PRRX1* was sufficient to increase the number of somatic LPM progenitors in the developing emu forelimb, we overexpressed *PRRX1* in both chicken and emu somatic LPM via targeted electroporation. Curiously, this was not sufficient to increase the number of somatic LPM progenitors in the early limb buds relative to the un-electroporated side (Figure S5). While *PRRX1* is a known lineage marker of the LPM and limbs, its role during limb development is unclear, where individual and *Prrx1/Prrx2* compound mouse mutants possess mild polydactyly but not overall effects on limb size ^50,51,54,55^. Instead, we looked at the origin of *PRRX1* activation, which has been observed through ectodermal BMP signalling ^11,16,56^. BMP signals from the overlying ectoderm specify somatic LPM fate ^11,16,56^ through induction of somatic LPM genes, including *PRRX1* and *TBX5* ^16^. Therefore, given the observed downregulation of both *PRRX1* and *TBX5* in the emu, it raised the question whether the reduced allocation of emu somatic LPM progenitors may arise through attenuated BMP signalling. We were unable to detect compelling differential expression of BMPs or BMPRs between the chicken and emu ectoderm and LPM, respectively (Table S2,S3). However, signalling pathway activity may be altered through the combined regulation and activity of ligands, receptors and/or SMAD transducers to influence altered outcomes. We therefore additionally predicted ligand-receptor interactions between the ectoderm and LPM in both chicken and emu and looked for the presence of differential signalling activity using CellChat, further failing to detect altered BMP pathway activity between the chicken and emu (data not shown). Therefore, while we detect a decreased allocation of limb progenitors within the somatic LPM (Figure 2) and downregulation of somatic LPM genes in emu (Figure 3), it is unclear whether this is cell autonomous or influenced by tissues within the surrounding limb field.

## Discussion

Of the extant flightless ratites, the emu has evolved the most vestigial wing reduction, both in size and number of digits ^6,37^. Previous work investigating the developmental basis of this reduction have identified reductions to several molecular mechanisms occurring after initiation of forelimb development ^35,36,38–41^. However, some evidence implied an earlier origin during development of the somatic LPM ^38,40^. Here, we resolve this question using comparative single cell transcriptomics and spatial quantification to analyse cell type proportions and disparate mechanisms underlying development of the emu and chicken embryonic forelimb field. We demonstrate that reduced emu wing development begins at the earliest origins of limb formation, via a reduced commitment of limb progenitor cells within the somatic LPM during formation of the presumptive forelimb field, known as heterotopy. This reduction is accompanied by downregulation of genes associated with LPM development and early limb specification ^16^, prior to activation of the canonical limb induction pathway which was previously implicated in driving vestigial emu wing development ^38,39^. These data demonstrate that stunted formation of the emu wing first occurs via a heterotopic reduction of forelimb progenitor cells, rather than a heterochronic delay in limb formation as previously thought ^38,40,42,43^. While the emu LPM arises at similar developmental stages to the chicken, the somatic LPM is reduced and contributes fewer TWIST1+ limb progenitors to the developing limb bud mesenchyme (Figure 2,3). This emphasizes that the evolution of divergent limb morphologies does not exclusively occur through post-patterning of the established limb bud primordia ^29–35^. Rather, these findings present an alternate mechanism whereby heterotopic differences in the allocation of limb progenitors can instead influence limb size, shape, or proportions within or between species (Figure 4) ^57^.

**Figure 4.**
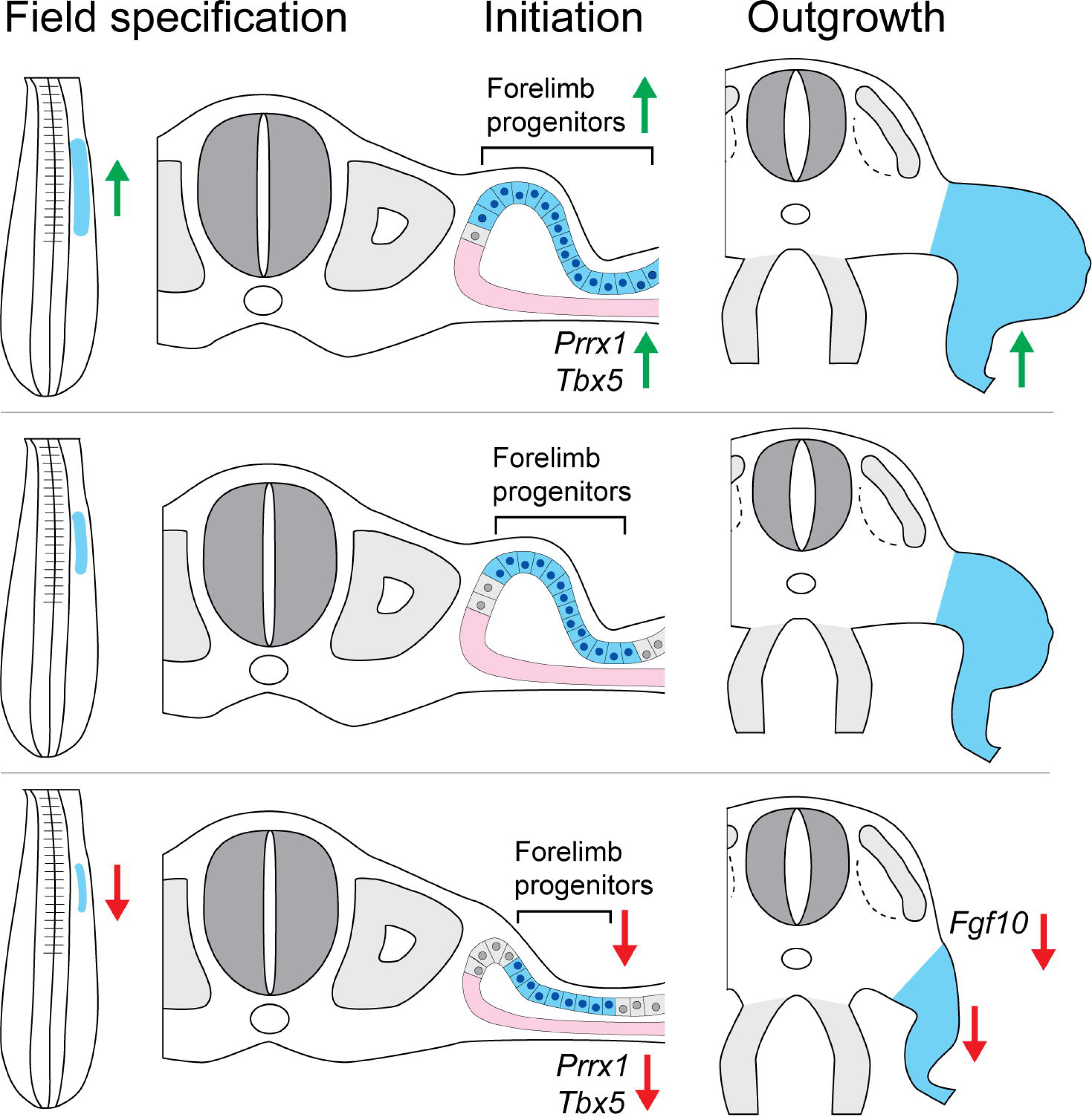
Progenitor allocation model during limb development. Altering the proportion or allocation of limb progenitor cells within the somatic LPM can influence the size and rate of growth of the developing limb. More progenitors (top; as seen in marsupials) results in enhanced limb gene expression and thus increased limb size; while fewer progenitors (bottom; as seen in emu) results in decreased gene expression and stunted limb formation.

While this is the first evidence for a role of heterotopy underlying differential limb development in avians, heterotopy has been shown to be a major contributor to enhanced forelimb development in marsupial embryos ^58^. Marsupials have evolved accelerated forelimb development to support the functional demands of crawling into the pouch by the altricial neonate ^58–62^. To achieve the rapid formation of large and well developed limbs, marsupials have evolved enhanced formation of somatic LPM-derived limb progenitors through a rostrocaudal-expanded forelimb field, compared with other mammals ^58^. This expansion ensures the marsupial embryo has a larger pool of somatic LPM and somite-derived limb progenitors to accelerate formation and outgrowth of the limb bud mesenchyme and musculature ^58,63,64^.

While we find that fewer progenitor cells are specified in the emu, the inductive signals which drive the reduced commitment of these remain unclear. Previous work has implicated BMP signalling from the overlying ectoderm as the inductive source of limb progenitor specification ^11,16,56^. However, we did not find any compelling evidence to suggest attenuated BMP signalling drives the reduced specification of limb progenitors in the emu, nor does ectopic overexpression of *PRRX1*, a target of BMP signalling, in the somatic LPM increase numbers of limb progenitor cells. This raises questions into how heterotopic differences in specification of limb progenitors is achieved, and whether this is cell-autonomous (genetically predisposed) or extrinsically influenced by surrounding tissues within the limb field. In the case of the former, it is plausible that the convergent cis-regulatory homoplasy observed amongst ratites ^8^ may indeed influence heterotopic reductions of progenitors across flightless species, acting as a shared mechanism underlying development of a vestigial wing. This hypothesis would benefit from additional observations of the size of somatic LPM and PRRX1/TWIST1+ limb progenitor fields across other ratite species. However, this does not appear to be the case in the ostrich ^12^, though compared to other ratites has retained large, semi-functional wings^9,35,43^. Conversely, while the adaptation of ratites towards a terrestrial environment has negated the need for functional wings, it has simultaneously increased requirements for large, robust hindlimbs. Relative to the chicken, the observable increase in spatial and temporal activation of *PRRX1* – demarcating future limb progenitor cells ^53,54^ in the developing emu hindlimb (Figure 3), coupled with upregulated expression of hindlimb genes seen in other studies ^38^, leads us to speculate whether the emu (and ratites more generally) may have additionally evolved a heterotopic increase of hindlimb progenitors to allow development of their robust hindlimbs.

Together, these observations lead us to propose a ‘progenitor allocation’ model whereby the development of different sized limbs may be initially patterned by changes to the proportions of the underlying limb progenitors (Figure 4). Not only does this present an acutely testable hypothesis that can be comparatively applied to different vertebrate species, but a broader evolutionary mechanism which underpins limb development between distantly related vertebrates. Furthermore, the progenitor allocation model may be further applicable to the ‘Allocation and Expansion’, or ‘Early Specification’ model of proximodistal limb patterning.

This hypothesis proposes that limb bud mesenchyme cells in the early limb ridge are initially allocated to future stylo, zeugo and autopod segments, before undergoing region-specific proliferation to form the limbs ^65,66^. It is intriguing whether these models may be intrinsically linked, helping to fill some gaps in knowledge about how early progenitor pools may arise ^67^. Altering the number of cells in somatic LPM-derived limb progenitor pool could influence the quantities allocated to formation of each proximodistal segment, thus influencing overall size. In the case of the emu forelimb, this may (partially) explain the overall size reduction of each of the segments, as well as the reduction of the typical avian autopod from three digits to one ^37^.

In this study we shed light on the intricate molecular mechanisms underlying vestigial wing development in the flightless emu. Our findings reveal that the emus stunted forelimb development occurs at the earliest stages of limb formation, through reduced specification of limb progenitors, rather than delayed development as previously thought^38,40,42,43^. Through comparative single-cell transcriptomics and spatial quantification, we demonstrate a heterotopic reduction in limb progenitor cells within the somatic LPM during the formation of the presumptive forelimb field. This early alteration in progenitor allocation, accompanied by downregulation of key developmental genes, provides a novel perspective on the evolutionary dynamics of limb development. Our observations pave the way for a ’progenitor allocation’ model, offering a testable hypothesis with broader implications for understanding limb development across diverse vertebrate species. As the role of heterotopy in morphogenesis gains prominence, it serves as a crucial framework for unravelling the intricate interplay of genes and cellular behaviours which drive adaptive outcomes throughout evolution.

## Methods

### Egg incubation, tissue collection, single cell sampling

Fertilized chicken and emu eggs were sourced from Research Poultry Farm (Victoria) and Emu Logic (NSW), respectively. Chicken sampling has been described previously ^16^, but was performed concurrently with emu egg collection. Eggs were incubated at 37.5°C using staggered timepoints so that different developmental stages were reached on the same day of collection. Chicken eggs were collected at embryonic day (e) 1.5 (stage 10), e2.5 (HH14) or e3.5 (HH18), while emu eggs were collected at e3.5 (HH10) e4.5 (HH14) and e5.5 (HH18). Embryos were dissected away from extra-embryonic membranes, rinsed in ice-cold DPBS then the LPM dissected. LPM tissues were digested with 0.05% Trypsin / EDTA and incubated at 37°C for 15 minutes, with mechanical dissociation every 5 minutes until no clumps were visible. Enzymatic activity was stopped with addition of 10% FCS. The dissociated cells were spun at 400g for 5 minutes, then resuspended in 1x EDTA / Propidium Iodide in DMEM (Gibco). Cells were filtered through a 70um Flowmi Cell Strainer (Scienceware), and viable cells were isolated through flow cytometry (Flowcore, Monash University). Chicken (n=3) and emu (n=3) samples were produced on different days and submitted to Micromon Genomics (Monash University) for independent processing through the 10X Genomics Chromium controller, using Single Cell 3’ V3.1 chemistry, as per the manufacturer’s instructions. Samples were subjected to 10 cycles of PCR for cDNA amplification and 16 cycles for library amplification. Once chicken and emu libraries had been constructed, chick they were sequenced using on the MGITech MGISEQ2000RS platform with MGIEasy V3 chemistry, on two lanes with 100bp paired end reads.

### Quantification and statistical analysis, bioinformatics

Chicken data analysis has been described previously ^16^, but was performed concurrently with emu analysis. Chicken reads were aligned to GRCg6a, and emu to droNov1 reference genomes using CellRanger (v4.0.0, using option: –force-cells 15000). Due to the number of reads observed just downstream of annotated genes, the gene annotation (from Ensembl release 100, gene biotypes: protein coding, lincRNA and antisense) was edited to include 1000bp downstream of each gene. Single cell analysis was performed in R using packages scran ^68^, scater ^69^ for QC, monocle3 ^49^ for trajectory analysis, and iSEE for interactive viewing ^70^. Gene names were used for analysis and, where they mapped to multiple Ensembl ids, the Ensembl ID with the highest number of counts was kept. Cells with low total UMI counts (<2000) were excluded. Cell cycle was annotated with cyclone in the scran package ^68^ using the mouse reference from ^71^ mapped to its one-to-one chicken orthologs.

For each dataset, the top 1000 genes with the highest biological variance were identified with modelGeneVar function of scran ^68^, blocked on the sequencing sample, and excluded mitochondrial genes or genes on the Z or W chromosomes to minimise sex effects. PCA was calculated on these, and the first 15 PCs used to generate a global chicken and emu tSNE layout. Clusters were defined with the walktrap method, on a SNN graph (k=10) ^68^, and cluster identities were determined from gene logFC changes and spatial expression profiles in the Gallus Expression *In Situ* Hybridization Analysis (GEISHA) database ^46,47^. Each dataset showed similar cells cluster labels, albeit with minor differences.

### Dataset integration and batch correction

To integrate the full chicken ^16^ and emu datasets, these were filtered to contain shared 1:1 gene IDs. Chicken and emu gene ID were intersected to produce a common ‘universe’ of shared features (n = 10415), and each dataset was subset to only contain these gene IDs and synchronized to contain the same metadata columns. Each dataset was rescaled to adjust for sequencing depth using batchelor, and integrated with Mutual Nearest Neighbours (MNN) correction ^48^ with k=20 neighbours and d=50 dimensions. This produced a corrected matrix of cell co-ordinates which was used as input for PCA, UMAP and tSNE dimension reduction, which showed good mixing of chicken and emu samples, recapitulating all of the known tissue types within the early forelimb field ^16^. Raw and log count assays and cell metadata was re-added into the corrected matrix, yielding a final merged and batch corrected dataset.

### Mesoderm analysis and lineage reconstruction

Mesodermal cell clusters were subset from the combined chicken and emu dataset for comparative analyses. Briefly, mesodermal clusters were subset to a new object, PCA and UMAP dimension reduction was recalculated using the top 250 highly variable genes and corrected for cell cycle effects. Next, the object was imported into Monocle3 ^49,72^ for k-means clustering (k=4). Cluster labels were confirmed by identifying differentially expressed marker genes through regression analysis implemented in monocle *fit_models* function and compared to those identified previously ^16^. This resulted in 9 sub-clusters covering the major cell types within the developing limb field mesoderm.

Lineage trajectories throughout the mesoderm were constructed in Monocle3 using reverse graph embedding (k=4, minimum branch length = 15, rann.k = 50), which produced 3 major lineages that originated in E1.5 cells and terminated in E3.5 cells. Lineage bifurcation points were corroborated using know LPM marker gene expression through the *plot_cells* function, then pseudotime was calculated by selecting the origin of the lineages using the *order_cells* function. Cell type proportions were investigated along lineage trajectories and visualized as relative between chicken and emu across each stage.

### Differential expression analysis

To perform differential expression between chicken and emu homologous clusters we used a pseudobulk approach. Cluster-specific cells were aggregated across sample, and samples with < 5 cells were removed. DE between chicken and emu clusters was performed using the *pseudoBulkDGE* function in *scran,* using stages as replicates. DE genes were visualised as volcano plots and through the *plot_cells* function in Monocle3. To identify genes that dynamically changed in expression across pseudotime, the LPM > somatic LPM > limb bud mesenchyme lineage was subset using the *chose_graph_segments* function in Monocle3. Pseudotime values for each cell were extracted and DE expression between chicken and emu genes along pseudotime were calculated using R packages ‘*splines’* and ‘*limma’*.

### Gene expression analysis by in situ hybridization and immunofluorescence

Whole mount *in situ* hybridization for spatial mRNA expression was carried out as described previously ^16^ with minor modifications. Briefly, whole HH8-HH22 chicken embryos were fixed overnight in 4% paraformaldehyde, dehydrated in methanol, and rehydrated in PBTX (PBS + 0.1% Triton X-1000). Tissues were permeabilized in 10µg/mL proteinase K for up to 1 hour, depending upon stage, then re-fixed in glutaraldehyde/ 4% PFA. Tissues underwent pre-hybridization (50% formamide, 5x SSC, 0.1% Triton X-100, 0.5% CHAPS, 5mM EDTA, 50mg/mL Heparin, 1mg/mL yeast RNA, 2% blocking powder) overnight at 65°C. The complete chicken *PRRX1* ORF was used as the riboprobe template, and amplified from limb and whole embryo cDNA using primers F-ATGGCGTCCAGCTATGCC and R-TTAATTGACTGTGGGCACTTG ^16^. Fragments were resolved by 1% agarose electrophoresis, excised, and purified using a Nucleospin PCR clean-up kit and subcloned into p-GEM T-easy (Promega). Antisense RNA probes were synthesized using SP6 RNA polymerase and the DIG-labelling kit (Roche, #11277073910) as per the manufacturer’s instructions. Precipitated probe was added to pre-hybridized tissues (approx. 5mL/ tube) and hybridization was carried out overnight at 65°C. Tissues were then subjected to stringency washes, blocked in BSA, then treated overnight with anti-DIG antibody conjugated with alkaline phosphatase. Tissues were exposed to BCIP/NBT colour reaction at room temperature for up to 3 hours (340mg/mL NBT and 175 mg/mL BCIP in NTMT (100mM NaCl, 100mM Tris-HCl, pH9.5, 50mM MgCl2, 0.1% Tween-20).

Chicken and emu embryos were fixed in 4% PFA/PBS for 15 minutes at room temperature then cryo-protected in 30% sucrose. Embryos were snap frozen in OCT and 10mm frozen sections were cut. Antigen retrieval was performed by the Monash Histology platform, where slides were heated at 60 degrees for 30 minutes followed by retrieval in citrate solution (pH 6). Sections were blocked and permeabilised in 1% Triton X-100, 2% BSA/PBS for 1-2hr at room temperature, then incubated with mouse anti-TWIST1 primary antibody (ab50887, Abcam) at 1:100 concentration in 0.5% Triton X-100, 1% BSA/PBS overnight at 4°C. An anti-mouse secondary (A21202, Invitrogen) was added at 1:1000 the following day and left at room temperature for one hour. Slides were washed once with DAPI diluted in PBS, followed by two washes in PBS.

## Acknowledgements

We thank Micromon Genomics for assistance with sequencing design, Monash University Flowcore for flow cytometry, and Monash Histology platform for histological processing. We additionally thank Nadia Davidson, Alex Combes and Kieran Short for helpful suggestions for choice of bioinformatics methods, and constructive comments from all members of the Smith Lab. This work was supported by the Australian Research Council Discovery Project scheme (DP190100890 to Craig Smith).

---

**Figure S1. Individual UMAP visualization and clustering of chicken and emu mesoderm.** Cell types were in agreement between the two datasets, show all mesoderm-derived cell types in the developing forelimb field. NKX2-5 has previously been noted to be co-opted in the developing emu forelimb ^41^, which we find expressed in early somatic LPM cells and the somites, which is not present in the chicken.

**Figure S2. Labelling of chicken mesoderm clusters on integrated chicken and emu dataset.** Clusters were largely in agreement between the chicken atlas ^16^ and merged data.

**Figure S3. Gene ontology enrichment of emu downregulated genes across pseudotime.** Enriched terms included forelimb morphogenesis and development.

**Figure S4. Spatial and temporal *in situ* visualization of *TBX5* gene expression during chicken and emu forelimb development.** This emu possessed reduced and delayed activation of *TBX5* in the developing forelimb field (black arrow) despite covering a similar number of somites, relative to the chicken. Note the shorter and narrower width of the *TBX5* positive limb progenitor field in the emu forelimb.

**Figure S5. Ectopic expression of *PRRX1* does not rescue emu limb outgrowth.** Overexpression of *CAG_Prrx1_T2A_GFP* plasmid in the emu and chicken LPM does not alter the number of limb progenitor cells, compared to unelectroporated control sides.

**Table S1. Chicken and emu cells sampled per stage.**

**Table S2. Global differential expressed genes in chicken cell clusters.**

**Table S3. Global differential expressed genes in emu cell clusters.**

**Table S4. Differentially expressed genes between chicken and emu across limb developmental pseudotime.**

