## Supplementary figures and images for "Heterotopic reduction of forelimb progenitors underpins development of the vestigial emu wing; implications for vertebrate limb evolution"

### Figure S1

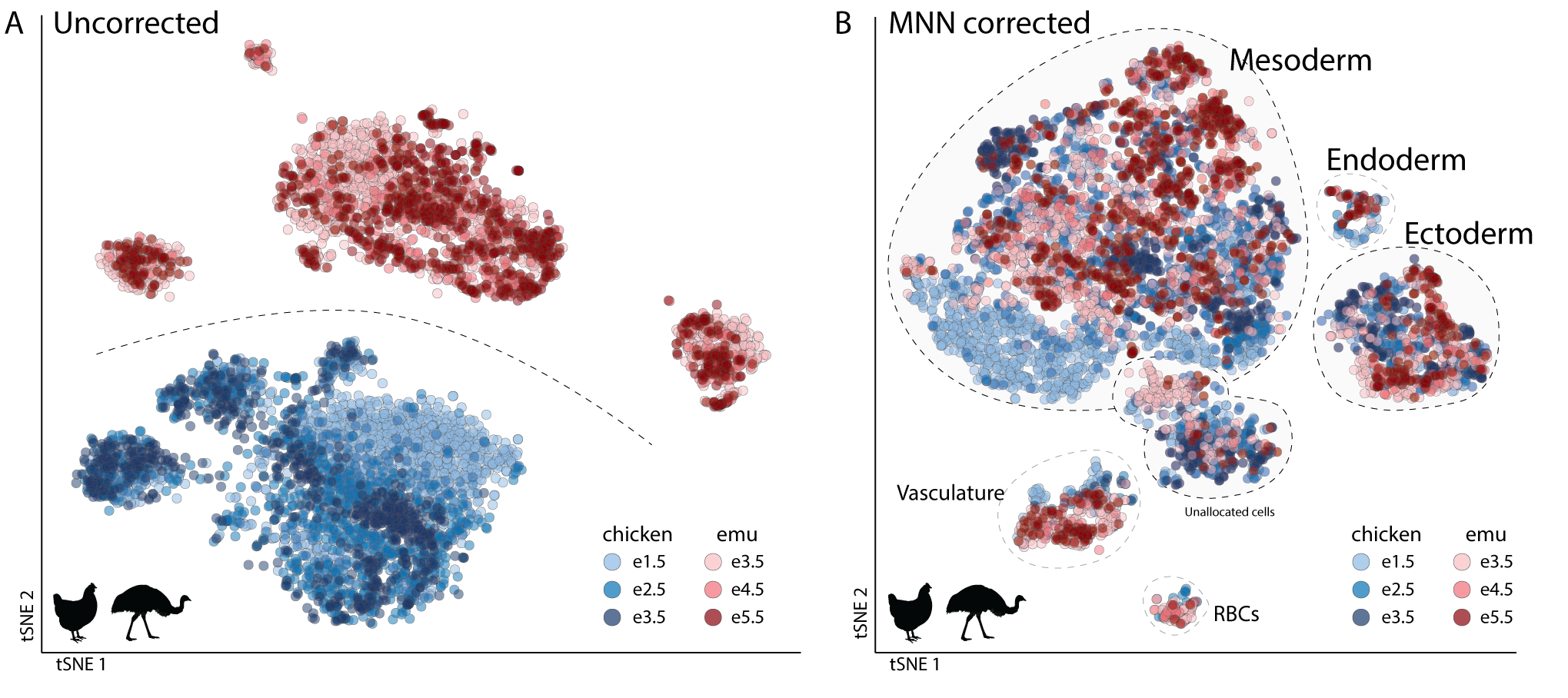

### Figure S2

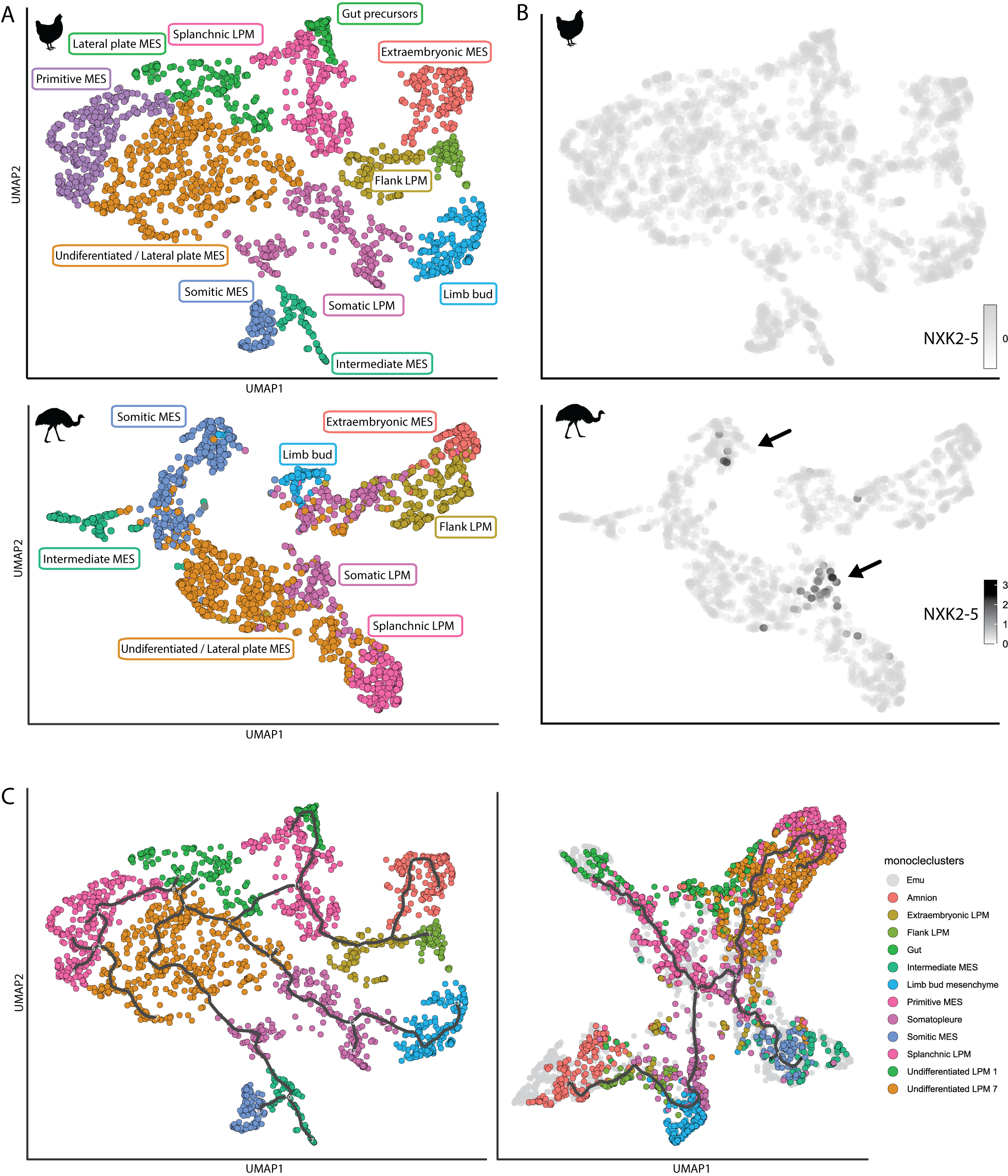

### Figure S3

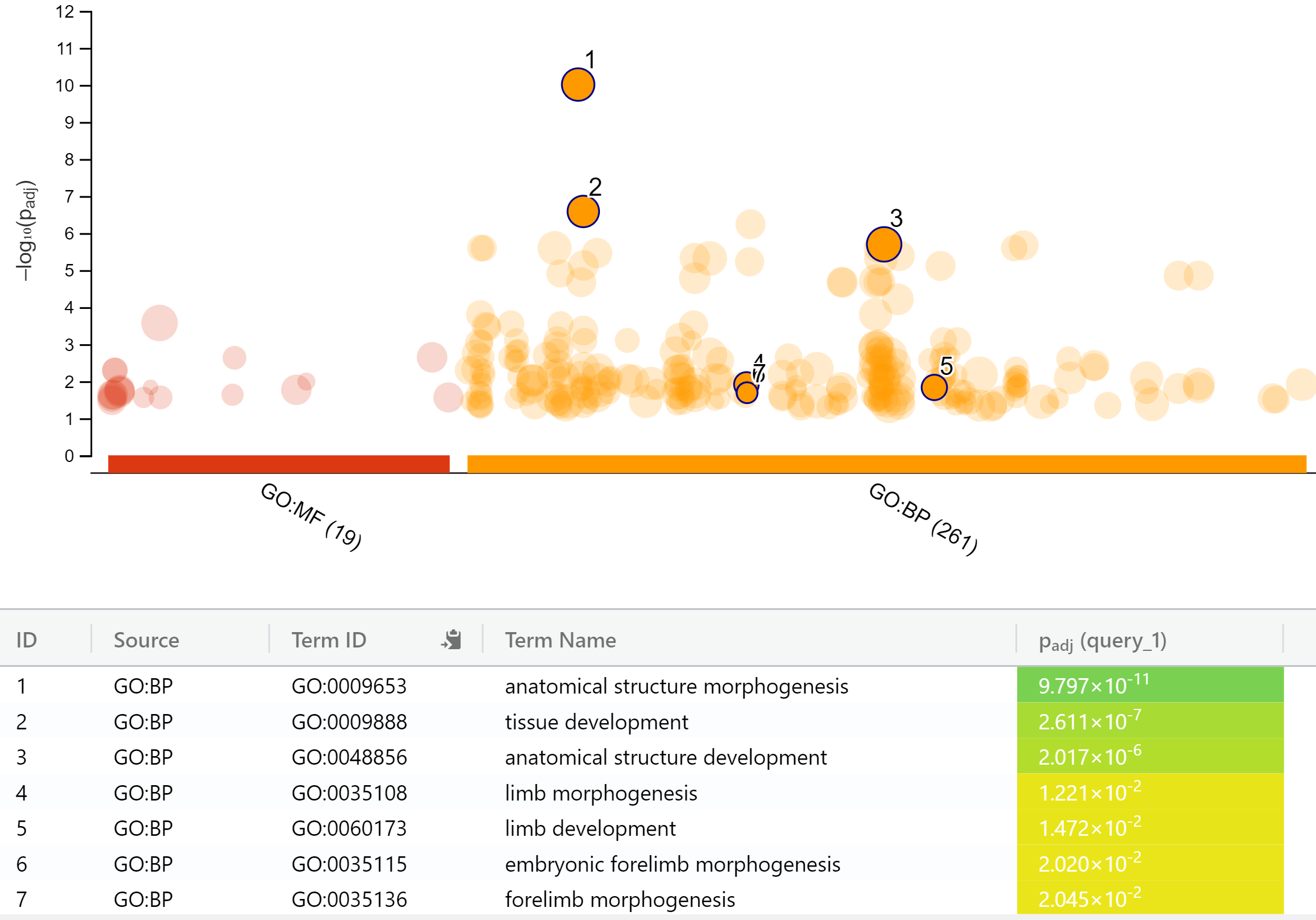

### Figure S4

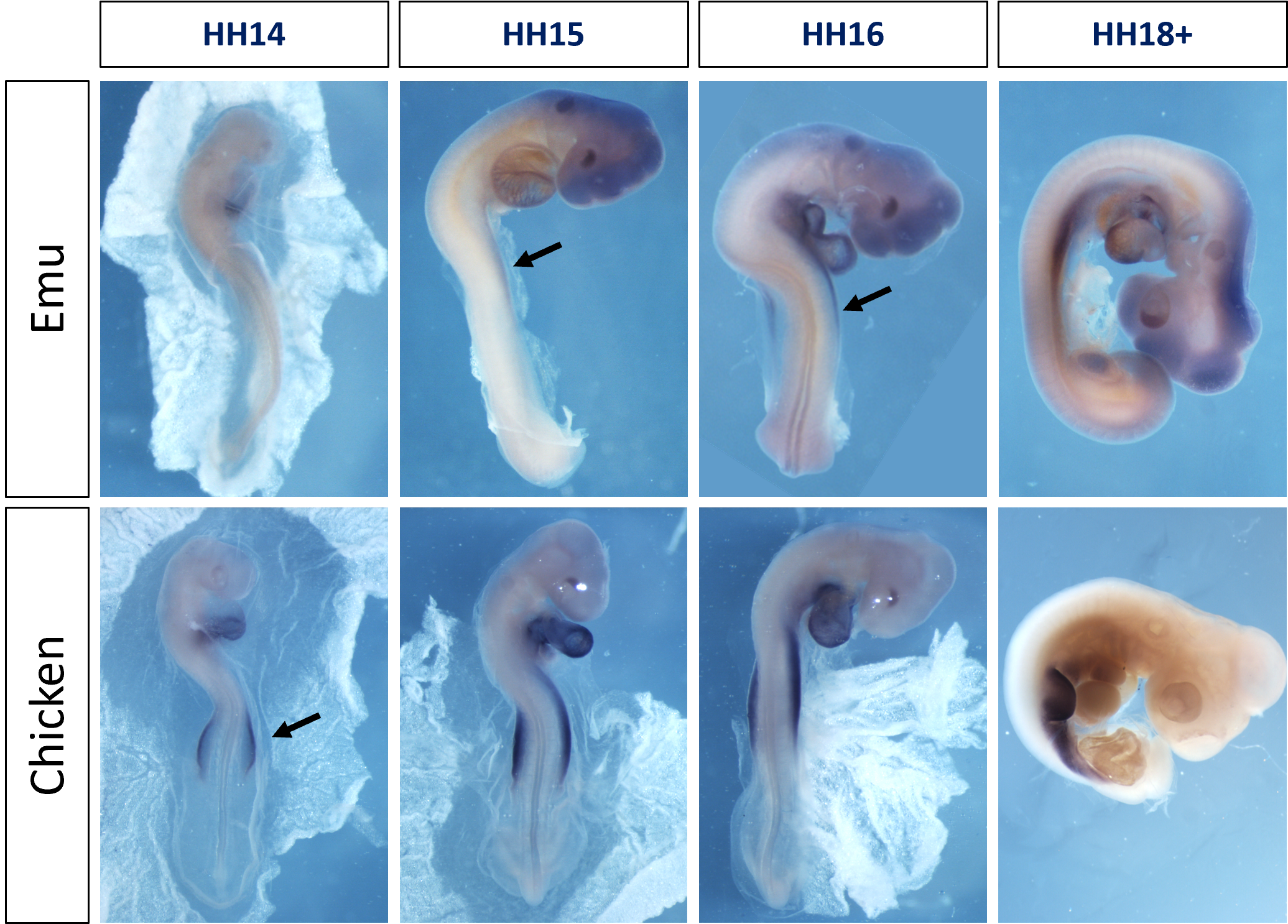

### Figure S5

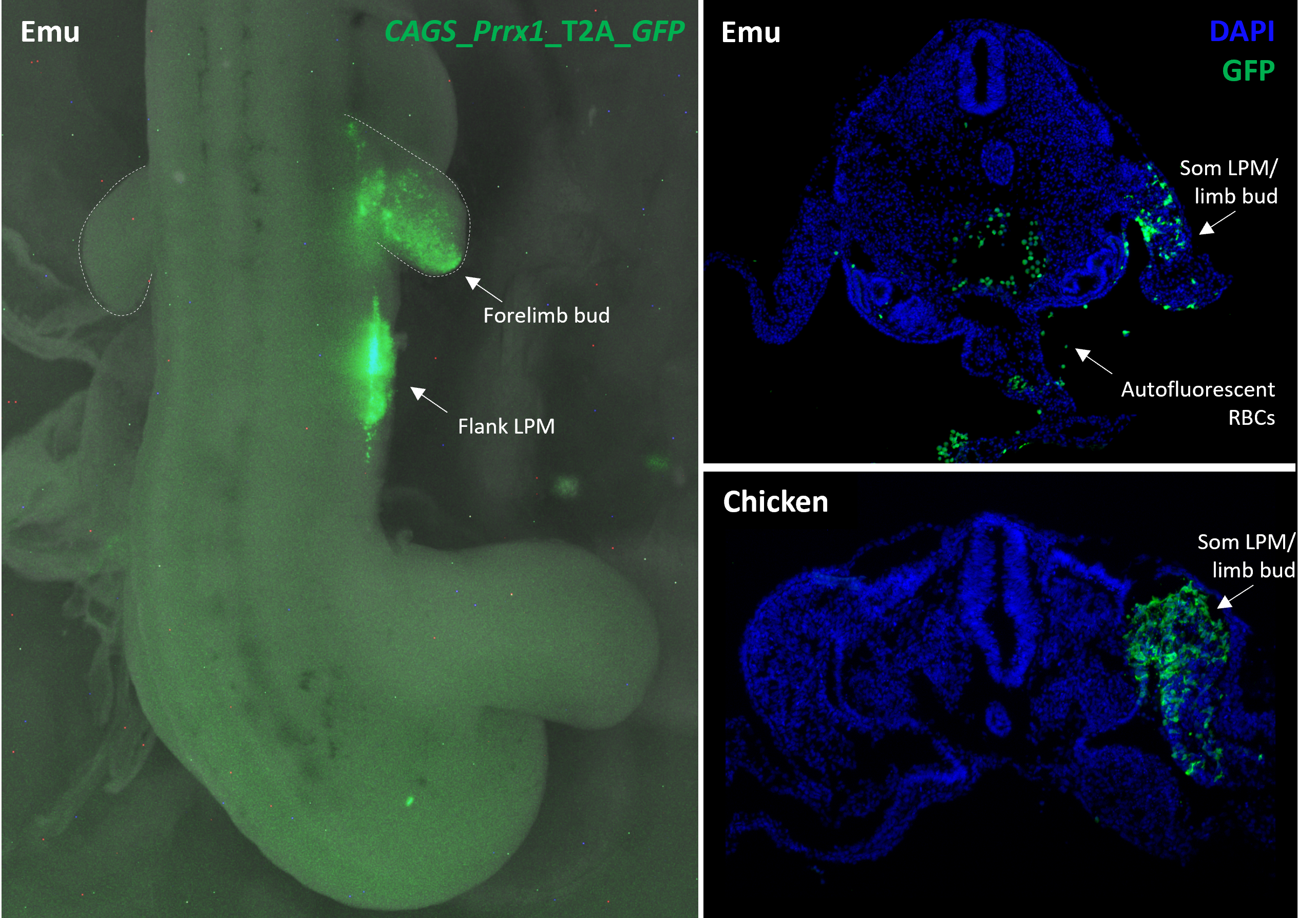
